# Rapamycin Differentially Impacts Germline Stem Cell Quiescence Across Diverse Genetic Backgrounds of *Drosophila Melanogaster*

**DOI:** 10.64898/2025.12.03.689791

**Authors:** Sahiti Peddibhotla, Miriam Gonzaga, Tricia Zhang, Yasha Goel, Jun Sun, Benjamin R. Harrison, Daniel E. L. Promislow, Hannele Ruohola-Baker

## Abstract

In response to ionizing radiation, stem cells enter a state of temporary quiescence, safeguarding the stem cell pool for tissue regeneration. Quiescence requires the inhibition of mTORC1, a kinase complex that promotes growth and suppresses autophagy, and cell cycle re-entry requires the reactivation of mTORC1. The pharmacological inhibition of mTORC1 in quiescent stem cells by rapamycin prevents cell cycle re-entry and subsequent tissue regeneration. It is well established that pharmacological responses can vary across genetic backgrounds, however, the extent to which genetic variation can impact the quiescence response to rapamycin is unclear. Here, we tested the sensitivity of stem cell quiescence to rapamycin in different genetic backgrounds within the *Drosophila* Genetic Reference Panel. These analyses revealed substantial variation across different genetic backgrounds, indicating that genetic variation can modulate drug-induced effects on stem cell dynamics. Our analyses suggest that mitophagy, rather than DNA damage response, is associated with the persistence of quiescence and delayed tissue regeneration by rapamycin. This work underscores the critical role of genetic background in determining drug efficacy, highlighting important implications for the therapeutic application of rapamycin.

## Introduction

Stem cells undergo asymmetric divisions to maintain tissue homeostasis and renew the stem cell pool. Under stress conditions, maintaining this stem cell pool is critical for effective post-injury regeneration [1, 2]. One of the most consequential stressors to cells is genotoxic stress, which induces DNA and organelle damage. Failure to repair these damages leads to the accumulation of dysfunctional organelles, misfolded proteins, and genomic instability, which can either trigger apoptosis, impair regeneration, or lead to tumorigenesis [3–8].

When exposed to ionizing radiation (IR), stem cells protect themselves from death and permanent damage by entering a transient state of proliferative dormancy, or quiescence, while their differentiated progeny undergo apoptosis [2, 9, 10]. Quiescence is essential to maintain cellular integrity. Following irradiation, *Drosophila* germline stem cells (GSCs) enter a quiescence which can only be exited upon the repair of DNA double-stranded breaks (DNA-DSB) and the autophagy of dysfunctional organelles [11–13]. DNA damage response across multiple organisms is directed by the FOXO transcription factor, which upregulates DNA repair proteins [14, 15]. In *Drosophila* GSCs, among the downstream factors of FOXO activation that induce cell cycle arrest, is the inhibition of the mechanistic target of rapamycin complex I (mTORC1) [11, 12]. The inhibition of mTORC1 promotes autophagy and mitophagy to remove damaged organelles [16–18]. Once cellular integrity is restored, mTORC1 reactivation promotes the switch from quiescence to post-injury proliferation and tissue regeneration.

The inhibition of mTORC1 is a general requirement of stem cell dormancy. Reduced mTORC1 signaling characterizes quiescence across multiple stem cell types in mammals and flies, including hematopoietic, neural, and satellite cells, and is observed in mammalian embryonic diapause and cancer stem cell dormancy [19–24]. By preventing mTORC1 inhibition, with a knockdown of the negative regulator Tsc1, irradiated *Drosophila* GSCs cannot enter quiescence [12]. In quiescent GSCs, treatment with the mTORC1 inhibitor rapamycin prolongs quiescence and prevents cell cycle re-entry [12]. By preventing cell cycle re-entry, rapamycin can reduce stem cell-based regeneration, but can also have therapeutic applications by preventing unwanted tissue growth, for instance, from dormant cancer stem cells [25–27]. Rapamycin is a promising intervention in mTORC1-driven diseases, such as cancer; however, little is understood about the pharmacogenetics and how its effects may depend on genetic background.

A few studies have demonstrated that the effects of rapamycin can vary among individuals across populations [28–30]. For instance, mice display sex differences in lifespan extension with rapamycin [28]. Two fly studies have demonstrated variation in multiple responses to rapamycin, including fecundity and development, using the *Drosophila* Genetics Reference Panel (DGRP) [29,30]. The DGRP is a collection of ∼200 genetically diverse wild-derived lab-inbred strains and is a powerful tool to assess the impact of genetic variation on drug response [31, 32]. Rapamycin delays development in *Drosophila*, such that larvae treated with rapamycin take longer to develop to the pupal stage [33, 34]. However, among the DGRP, there is extensive variation in the sensitivity of development time to rapamycin, including some lines showing greater sensitivity than the common lab strain *w*^1118^, and others remaining resistant to doses over two orders of magnitude higher than the screening dosage [30]. Variation in sensitivity to rapamycin during pupation delay is a strong indication that *Drosophila* populations maintain standing variation that influences the response of rapamycin on mTORC1 effector pathways.

We have previously shown that rapamycin extends irradiation-induced quiescence in the *w*^1118^ background [11]. Given recent studies that demonstrate genetic variation in response to rapamycin, we postulate that the sensitivity of IR-induced quiescence to rapamycin might depend on genetic background. Here, we survey 10 DGRP lines, and *w*^1118^, for sensitivity to rapamycin in prolonging quiescence after irradiation. We first find that the effect of rapamycin on GSC quiescence depends strongly on genetic background. We then hypothesize that the variation in the quiescence response might be the same pharmacogenetic variation in the effect of rapamycin on developmental delay. If this is true, we predict that the degree of sensitivity in quiescence across these lines would correlate to the variation in their sensitivity during development. We instead find that sensitivity in pupation delay does not correspond to sensitivity in quiescence, suggesting that genetic variation in mTORC1 effector pathways is specific to cellular processes that are involved in development or in GSC quiescence. We then investigate the mechanism behind variation in sensitivity to rapamycin in extending quiescence by considering the two pathways that are required for the exit out of quiescence: DNA repair and mitophagy. We find that lines at extreme ends of sensitivity to rapamycin in quiescence show no differences in DNA damage repair following IR-induction. However, we find that they differ substantially in the time course of mitophagy. By using lines that vary in sensitivity to rapamycin, our study suggests that genetic variation in the regulation of mitophagy could explain variation in sensitivity to rapamycin in quiescence.

## Materials and Methods

### Fly Stocks and Culture Conditions

w^1118^ and DGRP stocks were obtained from the Bloomington Drosophila Stock Center at Indiana University. The following stocks used were: w^1118^ (RRID:BDSC_3605), DGRP-57 (RRID:BDSC_29652), DGRP-307 (RRID:BDSC_25179), DGRP-383 (RRID:BDSC_28190), DGRP-441 (RRID:BDSC_28198), DGRP-517 (RRID:BDSC_25197), DGRP-287 (RRID:BDSC_28165), DGRP-348 (RRID:BDSC_55019), DGRP-443 (RRID:BDSC_28199), DGRP-776 (RRID:BDSC_28229), and DGRP-712 (RRID:BDSC_25201).

### Gamma-Irradiation Treatment and Dissection

Two days prior to ionizing radiation treatment, flies (0-3 days old) were cultured in empty vials with standard yeast paste (active dry yeast in dH2O) for 48 hours at 25°C. On the day of irradiation, five unirradiated females from each genotype were dissected (within one hour of irradiation), and the remainder were transferred into empty vials without yeast paste and treated with 50 Gy of gamma-irradiation. A Cs-137 Mark I Irradiator was used to administer the proper irradiation dosage as instructed by the dosage chart. At every timepoint, five of the irradiated females per treatment and genotype were dissected, while the remaining irradiated females were flipped into new vials with the respective treatment in yeast paste with young unirradiated males added at a ∼1:1 ratio to females.

### Development Assay

For each DGRP line assayed, 150-200 parents aged 3-5 days laid eggs for 3 h at 25°C in egg chambers containing grape agar plates with yeast paste dissolved with either 20 uM rapamycin in 5% EtOH or solvent control (5% EtOH). Plates with embryos were collected after 3 h. Yeast paste was changed to standard yeast paste at 36 h post-oviposition, and larvae were measured at 72 h post-oviposition.

### Rapamycin Treatment

A 4 mM stock was prepared in 200-Proof ethanol and stored at -20°C From 4 mM rapamycin, 1 mL of 200 µM in 5% EtOH was prepared and mixed with 0.5 g of active dry yeast. To prepare control food, 1 mL of 5% EtOH was prepared in dH2O and mixed in 0.5 g of active dry yeast. For irradiated flies, treatment or control was administered to the side of vials for 48 hours. For larvae, 20 µM rapamycin treatment or solvent control in yeast paste was administered at the center of grape agar plates in the egg collection chamber for the duration of oviposition (3 h) and 36 h post-oviposition. Standard and supplementary yeast paste was prepared with 0.5 g of active dry yeast for every 1 mL of dH2O.

### Immunocytochemistry

Fly ovaries were dissected in cold PBS, immediately fixed in 4% paraformaldehyde for 15 minutes, washed in PBT (PBS containing 0.2% Triton X-100) three times for 10 minutes each, and stored in PBS at 4°C for 24-96 hours. Dissected samples from all timepoints were simultaneously blocked in PBTB (PBT containing 0.2% BSA, 5% normal goat serum) for one hour at room temperature. Ovaries were incubated overnight at 4°C with the following primary antibodies: mouse anti-1B1 (RRID: AB_528070 1:30), mouse anti-ɑ-spectrin (RRID mouse anti-Lamin C (RRID: AB_528339 1:30), rabbit anti-γH2AV (RRID: AB_828383 1:200). After three 10-minute washes with PBT, secondary fluorescence antibodies were utilized including anti-rabbit Alexa 488 (RRID: AB_221544 1:250) and anti-mouse 568 (RRID: AB_2535773 1:250) for 2 hours at room temperature. Samples were washed once with PBT for 10 minutes, incubated with DAPI (diluted with PBS to 2 μg/ml) for 15 minutes, and washed two times with PBS for 10 minutes each. The samples were mounted in glycerol and analyzed and imaged on a Leica SPE5 confocal laser-scanning microscope. Image processing and editing were performed on ImageJ. For all images, intensity was uniformly adjusted, and background was removed using ImageJ. Unedited images were used for γH2AV intensity quantifications.

### Intensity Quantifications

For every GSC in each treatment, line, and timepoint, the mean intensity over was calculated from ROIs drawn over γH2AV puncta within nuclei in ImageJ. Intensities of GSCs (n=10-19) were averaged for each timepoint and treatment, then normalized to the intensity of the untreated, unirradiated timepoint for each respective line. Normalized intensities were categorized into low damage (>0.5), moderate damage (0.5-2), and high damage (2<).

### Statistical Analysis

Student’s T test (two-way, unpaired), two-way ANOVA and Spearman’s correlation coefficient were computed using R. Graphs and diagrams were created using R, GraphPad Prism 10, and BioRender.

## Results

### Rapamycin Delays Post-IR Regeneration of the Drosophila Germline

We first aimed to determine the extent to which post-IR regeneration of the Drosophila germline is affected by rapamycin treatment. The Drosophila ovary is composed of approximately 16 developing follicles, or ovarioles, with six to seven developing egg chambers [35]. The germarium, typically housing two to three GSCs, sits at the anterior tip of the ovariole. The GSCs are adjacent to the stem cell niche, consisting of capsule cells (CpCs). Both the niche CpCs and apical terminal filaments (TF) can be identified with lamin C (LamC) to stain their nuclear lamina (Figure 1A). GSCs undergo asymmetric divisions to produce both a self-renewing stem cell and a daughter cystoblast (CB) [36].

**Figure 1.**
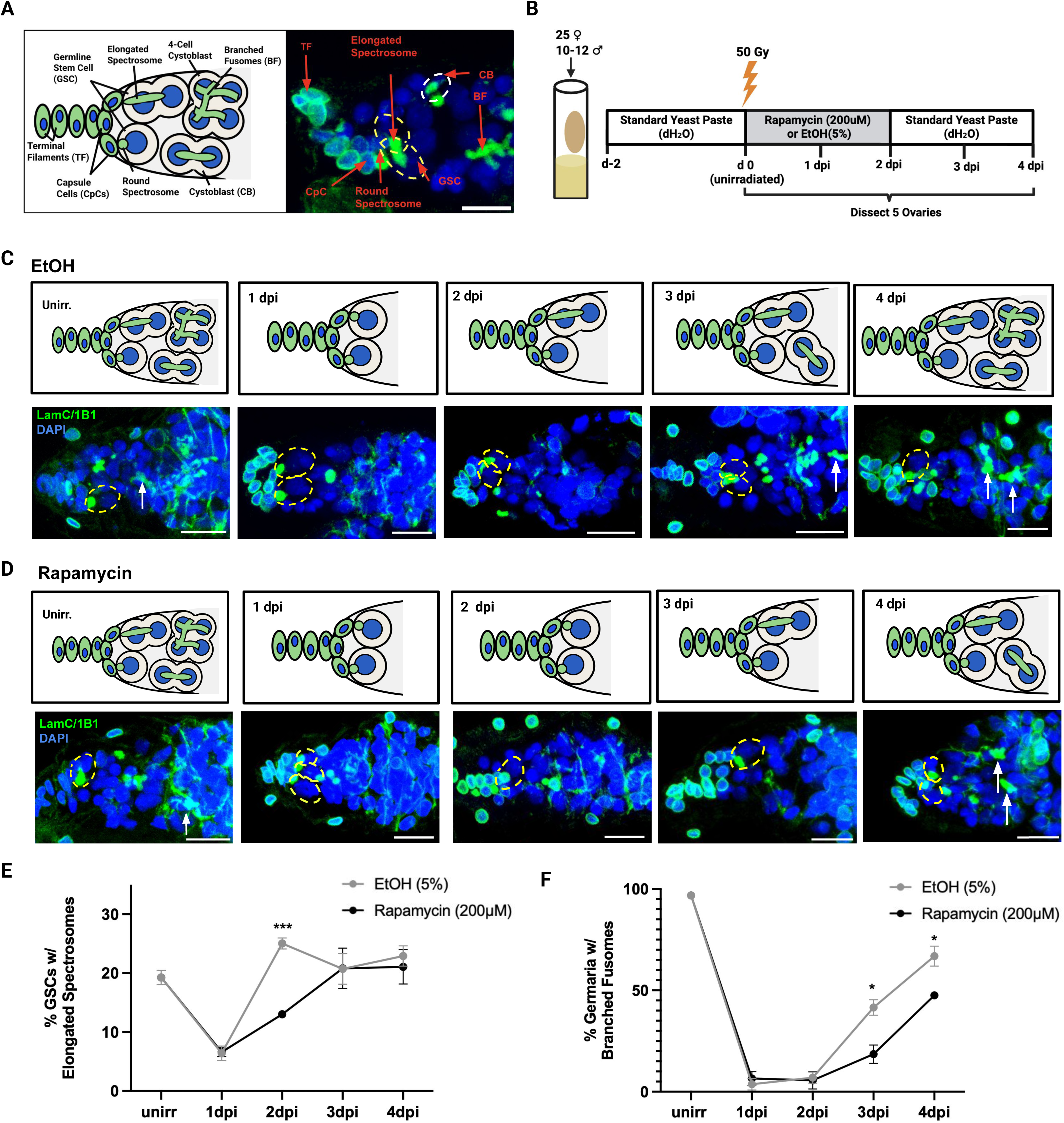
Rapamycin delays post-IR quiescence and regeneration in the Drosophila germline. (**A**) Representative illustration and image of an unirradiated w^1118^ control germarium, indicating the germline stem cell (GSC), daughter cystoblast (CB), and branched fusomes (BF). Terminal filaments (TF) are at the anterior-most (left) region of the germarium. Directly posterior to the TF are capsule cells (CpCs) that coat the germarium body. The GSCs are in contact with the CpCs, residing immediately posterior to them (Scale bar = 10 µm). (**B**) Experimental setup for irradiation (50 Gy) and rapamycin (200 µM) or solvent control (5% EtOH) treatment model. (**C**, **D**) Representative confocal microscopy images of germaria from w^1118^ flies treated with either solvent control (5% EtOH, C), or rapamycin (200 µM) in 5% EtOH (D). Germaria are oriented with the anterior facing left in all images. In C and D, germaria dissected from flies at unirradiated, 1 dpi, 2 dpi, 3 dpi, and 4 dpi timepoints were stained with LamC (green, CpC and TF), 1B1 (green, spectrosomes and BF), and DAPI (blue, nuclei). Spectrosomes are identified with 1B1 and are classified as round or elongated. GSCs are indicated by dotted yellow ellipses and fusomes are indicated by white arrows (Scale bar = 10 µm). (**E**) The percentage of GSCs with elongated spectrosomes in w^1118^ across all timepoints. Data represent mean ± SD from 147-253 GSCs across 3 biological replicates of 3-5 flies. (**F**) Percentage of germaria with BF in w^1118^ across all timepoints. Data represent mean ± SD from 117–212 germaria across 3 biological replicates of 3-5 flies. For (E-F), statistical significance was determined by two-way ANOVA (*p < 0.05, **p < 0.01, ***p < 0.001).

The cell cycle of the GSC can be tracked by examining spectrosome morphology, identified with spectrosome-specific antigens, such as 1B1 and spectrins. The S-phase of the cell cycle is characterized by a fused or elongated spectrosome which bridges a GSC and a CB. In Drosophila GSCs, injury-induced quiescence occurs at the G1/S transition. Elongated spectrosomes indicate that GSCs have re-entered the S-phase of the cell cycle and are no longer quiescent [12, 36]. The recovery of the germline follows GSC division and is accomplished when the daughter CB undergoes several rounds of incomplete divisions to produce a 16-cell cyst. One of the 16 cells then differentiates as the oocyte while the remaining cells differentiate as nurse cells to support the oocyte by synthesizing proteins and nutrients [37, 38]. As these incomplete divisions occur, CB spectrosomes branch into fusome networks, also marked by 1B1 and spectrins, whose presence indicate regeneration of the germline after IR. Following IR, the CB population is temporarily depleted but typically recovers fully by four days post-IR (dpi) [11, 15].

To assess the impact of rapamycin on germline regeneration, wildtype (w^1118^) flies (0-3 days old) were conditioned in food vials, supplemented with standard yeast paste, for 48 hours. Flies were then irradiated with 50 Gy to induce quiescence. Following IR, we treated flies with either 200 µM rapamycin in 5% EtOH or the solvent control (5% EtOH), dissolved in yeast paste, and analyzed GSC division rates by assessing GSC spectrosome elongation once every 24 hours. After 2 dpi, the treatment yeast paste (rapamycin or solvent control) was replaced with standard yeast paste in order to measure germline recovery by 4 dpi (Figure 1B) (Methods). In untreated flies, GSCs enter quiescence at approximately 1 dpi and then exit quiescence at approximately 2 dpi [11,12]. In germaria of both rapamycin-treated and control-treated flies, the percentage of GSCs with elongated spectrosomes significantly decreased at 1 dpi compared to the unirradiated (unirr.) timepoint, indicating successful induction of quiescence by IR (Figure 1C-E). At 2 dpi, the GSCs of rapamycin-treated flies continued to have a significantly lower percentage of elongated spectrosomes, compared to the GSCs of control flies at 2 dpi (Figure 1C-E). At 3 dpi, the percentage of GSCs with elongated spectrosomes in the rapamycin-treated flies had returned to a frequency similar to the unirradiated control samples (Figure 1E). Consistent with previous reports [11], we show that rapamycin delays quiescence in GSCs, preventing their re-entry into the cell cycle in the wild-type strain, w^1118^ (Figure 1E).

Next, we quantified the percentage of germaria containing BF (branched fusomes) as an indication of cystoblast division and germline regeneration. The percentage of germaria containing BF drastically decreases from unirradiated to 1 dpi, indicating death of somatic cells in the germline as a result of IR (Figure 1C-D, F). After 1 dpi, and until 4 dpi, BF frequency remained low in rapamycin-treated flies. In comparison, the BF frequency of control-treated flies returned to the value prior to irradiation more quickly (Figures 1C and 1F). At 4 dpi, 66.8% of germaria in control-treated flies contained BF, while a significantly lower percentage of germaria (47.5%) from rapamycin-treated flies contained BF (p = 0.02). This data indicates that rapamycin significantly delays post-IR regeneration of the germline.

### The Effect of Rapamycin Varies Across Genetic Backgrounds

While rapamycin can delay post-IR germline regeneration in the w^1118^ genetic background, which is commonly used in Drosophila melanogaster studies, it is unknown whether this effect is conserved among genetically diverse Drosophila strains. To test the initial hypothesis that rapamycin sensitivity in development corresponds to sensitivity in quiescence, we selected DGRP lines at the extremes of the sensitivity distribution reported in Harrison et al. (2024) (Figure 2A) [30]. The common lab strain, w^1118^, shows moderate sensitivity to rapamycin, delaying pupation by an average of 0.86 days, whereas more-sensitive strains show up to a 6-day delay by rapamycin, and resistant strains show little-to-no detectable delay (Figure 2A) [30].

**Figure 2.**
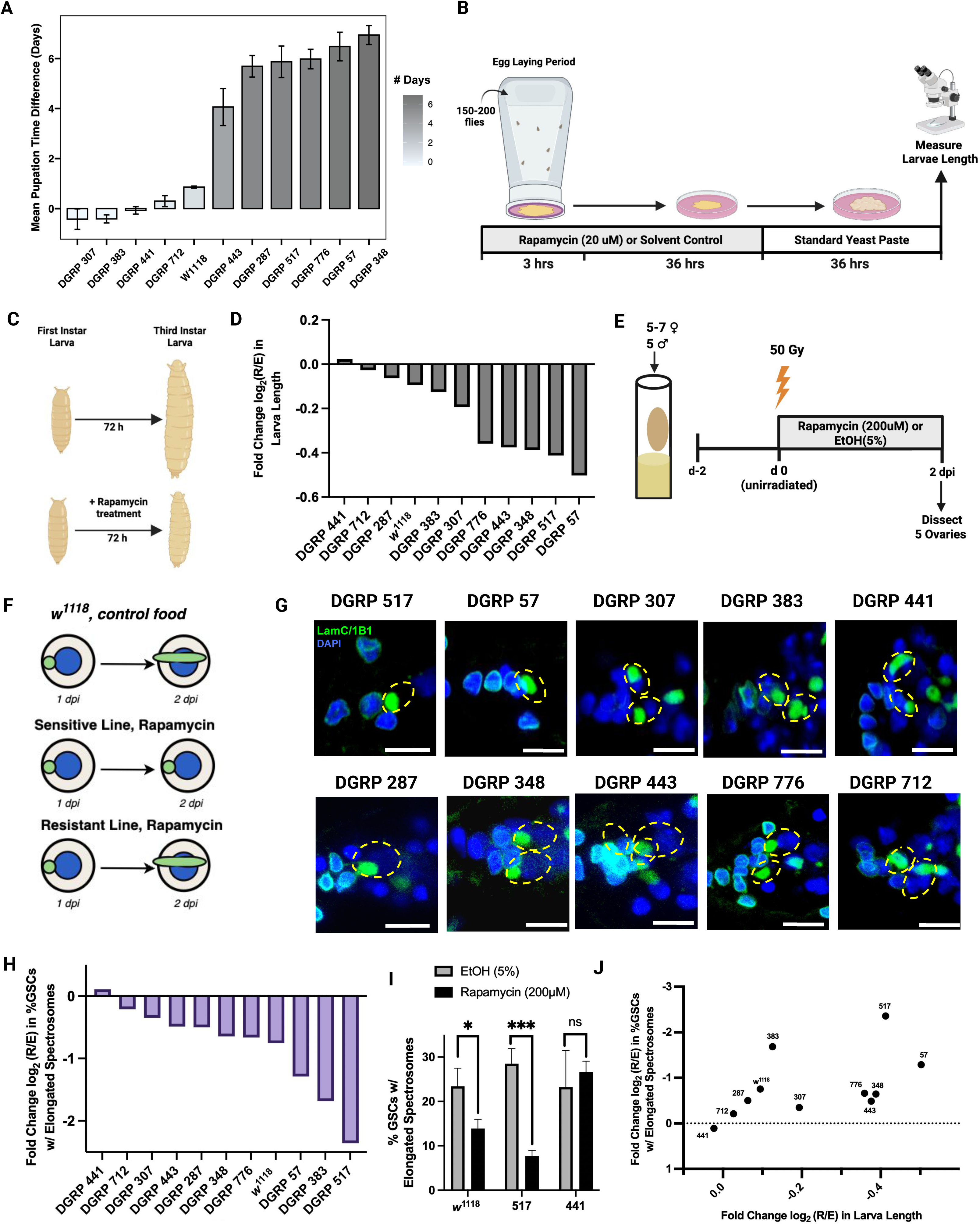
DGRP lines display variable response to rapamycin in larvae development and GSC quiescence. (A) DGRP lines showing extreme phenotypes of sensitivity and w^1118^. Data represents mean delay in pupation time between rapamycin and solvent control (days for pupation with control - days for pupation with rapamycin). Data acquired from Harrison et al. (2024). (**B**) Experimental paradigm for screening sensitivity to rapamycin in larvae growth. For each line and treatment, 150-200 parents (3-4 days old) were placed in chambers containing grape-agar plates with yeast diluted with solvent control (5% EtOH) or rapamycin (20 µM) in 5% EtOH. Eggs were collected after 3 hours. At 36 hours post-oviposition, larvae were transferred to new grape agar plates with standard yeast paste and measured after 36 additional hours. (**C**) Illustration comparing the change in size of sensitive larvae treated with rapamycin to control-treated larvae over 72 hrs. (**D**) Sensitivity to rapamycin in larvae length across w^1118^ and DGRP lines, as a log2 fold change in length (rapamycin/control) (n = 19-62 larvae). (**E**) Experimental paradigm for screening sensitivity to rapamycin in quiescence exit. 5-7 female flies from each DGRP line were irradiated (50 Gy), treated with either rapamycin (200 µM) in 5% EtOH or solvent control (5% EtOH) mixed in yeast paste, and dissected for ovaries at 2 dpi. (**F**) Illustration comparing the spectrosome elongation of GSCs from w^1118^ with control food and GSCs from rapamycin-treated lines that are sensitive or resistant to rapamycin across unirradiated, 1 dpi, and 2 dpi timepoints. (**G**) Confocal microscopy images of GSCs from each line, treated with rapamycin. GSCs were stained with LamC (green, CapC and TF), 1B1 (green, spectrosomes), and DAPI (blue, nuclei) (Scale bar = 5 µM). Germaria are oriented with the anterior facing left in all images. (**H**) Sensitivity to rapamycin in quiescence exit across w^1118^ and DGRP lines, as log2 fold change in spectrosome elongation rate (rapamycin/control) (n = 30-87 GSCs). (**I**) Spectrosome elongation data for lines w^1118^ (p = 0.023), DGRP 517 (p-value = 0.001), and DGRP 441 (p-value = 0.529). Significance calculated with Student’s T-Test (*p < 0.05, **p < 0.01, ***p < 0.001). (**J**) Scatterplot demonstrating the relationship between sensitivity to rapamycin in larvae length and sensitivity to rapamycin quiescence. Both length and quiescence are represented as a log2 fold change (rapamycin/control). Spearman coefficient (r_s_= 0.682, p-value = 0.021).

We reproduced the developmental phenotype of lines screened in Harrison et al. (2024) by measuring the length of 72 h old larvae, treated with 20 µM rapamycin in 5% EtOH or solvent control (5% EtOH) (Figure 2B, Methods). We recorded the phenotype of w^1118^, six lines that were identified as highly sensitive to rapamycin (DGRP 443, 287, 517, 776, 57, and 348), and four strains whose development was relatively resistant (DGRP 307, 383, 441, and 712). Drosophila larvae enter the early third instar stage at about 72 hours post-fertilization [39]. Early administration of rapamycin in Drosophila leads to reduced body size of third instar stage larvae (Figure 2C) [30, 40]. We find that the sensitivity to rapamycin greatly varies across the DGRP lines, where rapamycin reduces length by up to half in sensitive strains, and by almost none in resistant strains (Figure 2D). Larval length varies significantly with genotype and treatment, and we detect a significant genotype × treatment interaction (two-way ANOVA, p < 0.001 for all factors). Across these lines, the reduction in larval size by rapamycin shows a moderately strong correlation with the previously published pupation delay (r_s_= -0.655, p = 0.029, Supplementary Figure S1B) [30].

Next, we asked whether genetic variation impacts the delay in germline regeneration by rapamycin. The length of the GSC quiescent period influences the recovery of the germline from injury. Therefore, we observed the extent to which exit from GSC quiescence was delayed. GSCs from w^1118^ flies re-enter the cell cycle at 2 dpi, while GSCs in rapamycin-treated w^1118^ flies do not fully re-enter the cell cycle until 3 dpi (Figure 1). Therefore, we recorded the percentage of GSCs with elongating spectrosomes at 2 dpi in each line treated with rapamycin or solvent control to gauge how genetic variation affects cell cycle re-entry delay by rapamycin. We compared the response of w^1118^ to the responses of the six lines whose development was highly sensitive to rapamycin and to the four lines whose development was relatively resistant (Figure 2A). Flies were conditioned in vials with standard food and supplementary yeast paste for 2 days prior to IR. Following IR and treatment with rapamycin or solvent control until 2 dpi, ovaries were dissected to observe spectrosome morphology (Figure 2E).

We observed that rapamycin delays stem cell division post-IR by 1 day in most lines. However, the magnitude of this delay varies greatly amongst the lines (Figure 2G-H). w^1118^ showed moderate sensitivity, with a log2 fold change in spectrosome elongation of -0.76, with a significant decrease in elongation at 2 dpi in rapamycin-treated flies compared to control-treated flies (p = 0.023). Several other lines exhibited fold changes similar to w^1118^, such as DGRP 776 (-0.66) and 348 (-0.65). DGRP 517, whose developmental timing is highly sensitive to rapamycin, had a low fold change of -2.36 (Figure 2H), with a highly significant decrease in elongation at 2 dpi in rapamycin-treated flies compared to the control (p = 0.001) (Figure 2I). DGRP 441, whose developmental timing is highly resistant to rapamycin, had a positive fold change of 0.11 (Figure 2H) with no significant difference in elongation between treatments at 2 dpi (p = 0.529) (Figure 2I).

Sensitivity to rapamycin in GSC quiescence shows a moderately strong positive correlation with the effect on larvae length (r_s_ = 0.682, p = 0.021) (Figure 2J). However, comparing the sensitivity of GSC quiescence to pupation delay reported in Harrison et al. (2024), the degree of sensitivity to rapamycin in pupation delay does not significantly correlate with sensitivity in quiescence exit (r_s_ = 0.4, p = 0.223, Supplementary Figure S2B). Among the lines that were the most sensitive in pupation delay (Figure 2A), the lines 287 (-0.5), 443 (-0.45), and 776 (-0.66) appeared relatively resistant in quiescence. DGRP 383, on the other hand, was resistant in pupation delay (Figure 2A), but was among the more-sensitive lines to rapamycin in the GSC quiescence response (fold change = -1.68, Figure 2H). We show that there is a significant, moderately strong correlation between the previously published pupation delay and larval length (Supplementary Figure S1B). However, the lack of correlation between pupation delay and quiescence exit suggests that the effect of rapamycin not only depends on genetic background, but also on the biological context.

### Germline Regeneration Varies with Rapamycin Across Genetic Backgrounds

We have shown that genetic variation influences the extension of quiescence by rapamycin in GSCs. Next, we asked if the variation in sensitivity to rapamycin in quiescence is also associated with variation in the capacity for post-injury regeneration. Among the lines screened for quiescence sensitivity previously, DGRP 517 was the most sensitive to rapamycin, while DGRP 441 was the most resistant (Figure 2H).

We first assayed how rapamycin impacts the capacity for post-injury regeneration in these two lines, along with w^1118^ for comparison, by tracking spectrosome elongation rates throughout the duration of recovery (until 4 dpi). Conditioning, irradiation, treatment, and dissection were performed and spectrosome elongation rates were quantified. At 1 dpi GSCs from both lines, regardless of rapamycin or control treatment, entered quiescence as evidenced by the drop in spectrosome elongation (Figure 3A-D). However, a higher percentage of GSCs had elongated spectrosomes in DGRP 441 at 1 dpi (12.9% and 17.9% for control and rapamycin, respectively) compared to DGRP 517 (6.0% and 3.1% for control and rapamycin, respectively), suggesting that IR is less likely to induce quiescence in the GSCs of DGRP 441 than DGRP 517.

**Figure 3.**
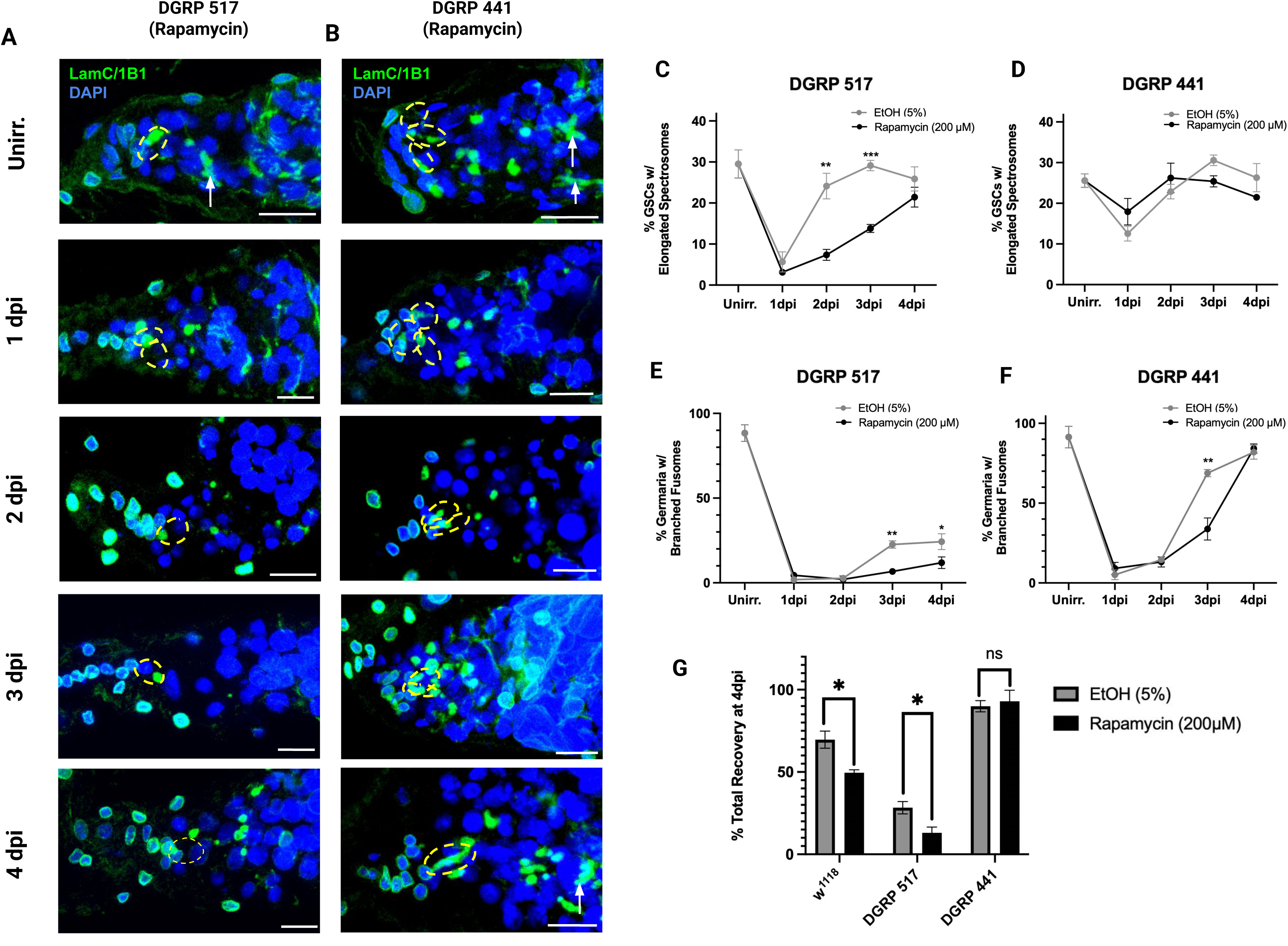
The germline regeneration of DGRP 517, but not DGRP 441, is sensitive to rapamycin. **(A, B)** Representative confocal microscopy images of germaria from DGRP 517 **(A)** or DGRP 441 **(B)** flies treated with rapamycin (200 µM) in 5% EtOH. Germaria are oriented with the anterior facing left in all images. In A and B, germaria dissected from flies at unirradiated, 1 dpi, 2 dpi, 3 dpi, and 4 dpi timepoints were stained with LamC (green, CpC and TF), 1B1 (green, spectrosomes and BF), and DAPI (blue, nuclei). Spectrosomes are identified with 1B1 and are classified as round or elongated. GSCs are indicated by dotted yellow ellipses and fusomes are indicated by white arrows (Scale bar = 10 µm). (**C**) Percentage of GSCs in DGRP 517 showing spectrosome elongation with solvent control (5% EtOH) or rapamycin (200 µM) in 5% EtOH (n= 76-166 GSCs). (**D**) Percentage of GSCs in DGRP 441 showing spectrosome elongation with solvent control (5% EtOH) or 200 µM rapamycin treatment (n=158-212 GSCs). (**E**) Percentage of germaria with BF in DGRP 517 with solvent control (5% EtOH) or 200 µM rapamycin treatment (n=71-154 germaria) (**F**) Percentage of germaria with BF in DGRP 441 with solvent control (5% EtOH) or 200 µM rapamycin treatment (n=101-131 germaria). Data for (C-F) represent mean ± SEM of 3 biological replicates of 3-5 flies. p-values were calculated using two-way ANOVA, *p < 0.05, **p < 0.01, ***p < 0.001. (**G**) Percent total recovery of the germline at 4 dpi as a percentage of germaria with BF normalized to the respective unirradiated line without treatment (% germaria with BF at 4 dpi/ % germaria with BF at unirradiated * 100%). Data represent mean ± SEM of n = 131-160 germaria from 3 biological replicates of 3-5 flies. P-values calculated with Student’s T-test (*p < 0.05, **p < 0.01, ***p < 0.001).

GSCs from DGRP 441 flies in both treatments exited quiescence by 2 dpi, as evidenced by the increase in spectrosome elongation (22.85% and 26.24% for control-treated and rapamycin-treated flies, respectively). In DGRP 441, there was no significant difference in percent GSCs with elongated spectrosomes between the treatments across all timepoints (Figure 3D). As observed formerly (Figure 2H, I), GSCs from DGRP 517 flies treated with rapamycin stayed in quiescence at 2 dpi while GSCs from those treated with the solvent control re-entered the cell cycle (Figure 3C). Unlike DGRP 441 and w^1118^, spectrosome elongation rates in rapamycin-treated DGRP 517 flies did not equate to those of the control-treated flies until 4 dpi (p = 0.308). We therefore affirm that the recovery of GSC division post-IR is compromised by rapamycin in the sensitive line.

Next, we assessed how regeneration of the germline varied in response to rapamycin by quantifying BF rates. BF rates in germaria from rapamycin-treated DGRP 441 flies were significantly lower at 3 dpi compared to control-treated flies, despite there being no significant difference in spectrosome elongation rates across all timepoints. However, by 4 dpi, the germline of DGRP 441 was fully regenerated, with 81.9% and 82.4% of germaria having BF for control-treated and rapamycin-treated flies, respectively, with no significant difference between the percent total recovery rates between treatments (p = 0.705). DGRP 517, on the other hand, showed a significant decrease in BF rates at 3 dpi and 4 dpi in the germaria of rapamycin-treated flies, compared to control-treated flies at those timepoints, with a significant difference in percent of total recovery at 4 dpi between the treatments (p = 0.041). Surprisingly, in both treatments of DGRP 517, the germaria do not fully recover despite there being high spectrosome elongation rates, suggesting that the cystoblast division does not fully recover after irradiation. Here we find that rapamycin significantly delays the post-IR germline regeneration of a sensitive line, but not a resistant one.

### Genetic Variation in Mitophagy and Not DNA Repair During Quiescence

We then test whether genetic variation in the response to rapamycin involves either of two repair mechanisms required to exit quiescence: DNA repair or mitophagy. Drosophila GSCs show evidence of DNA damage at 30 minutes post-IR which is fully repaired by 24 hours post-IR [11]. We tested whether the timing of IR-induced DNA-DSB repair differs between the highly sensitive DGRP 517 line and the highly resistant DGRP 441 line when treated with rapamycin, and in comparison, to the moderately sensitive genotype w^1118^. If rapamycin extends quiescence by impairing DNA damage repair, we should see that the DNA damage accumulation pattern of each line with rapamycin parallels the duration of quiescence.

To estimate the level of DNA damage, we measured the intensity of phospho-H2Av (anti-γH2Av), a marker for DNA-DSB in Drosophila (Methods). All lines and treatments showed an increase in γH2Av intensity from their unirradiated condition to 30 min post-IR (30 mpi, Figure 4A, B). In DGRP 517 and 441, both unirradiated and irradiated flies had a higher percentage of GSCs with high-level DNA damage compared to w^1118^, indicating that both lines have a higher sensitivity to IR (Figure 4B, D-E). In control-treated w^1118^ flies, all GSCs at 1 dpi had low-level γH2Av, suggesting they had repaired their DNA. In contrast, at 1 dpi in rapamycin-treated w^1118^ flies, 32% of GSCs still had moderate-level DNA damage. At 1 dpi, 91% and 80% of GSCs in control-treated DGRP 517 and 441 flies, respectively, showed moderate DNA damage. In the same lines, GSCs from rapamycin-treated flies at 1 dpi show similar percentages of moderate DNA damage (88% for DGRP 517 and 75% for DGRP 441), suggesting that rapamycin does not have a substantial effect on DNA repair in these lines. The similarity in DNA damage between rapamycin-treated and control flies of DGRP 517 and 441 suggests that DNA damage repair does not contribute to the substantial difference in the duration of quiescence in response to rapamycin in these lines.

**Figure 4.**
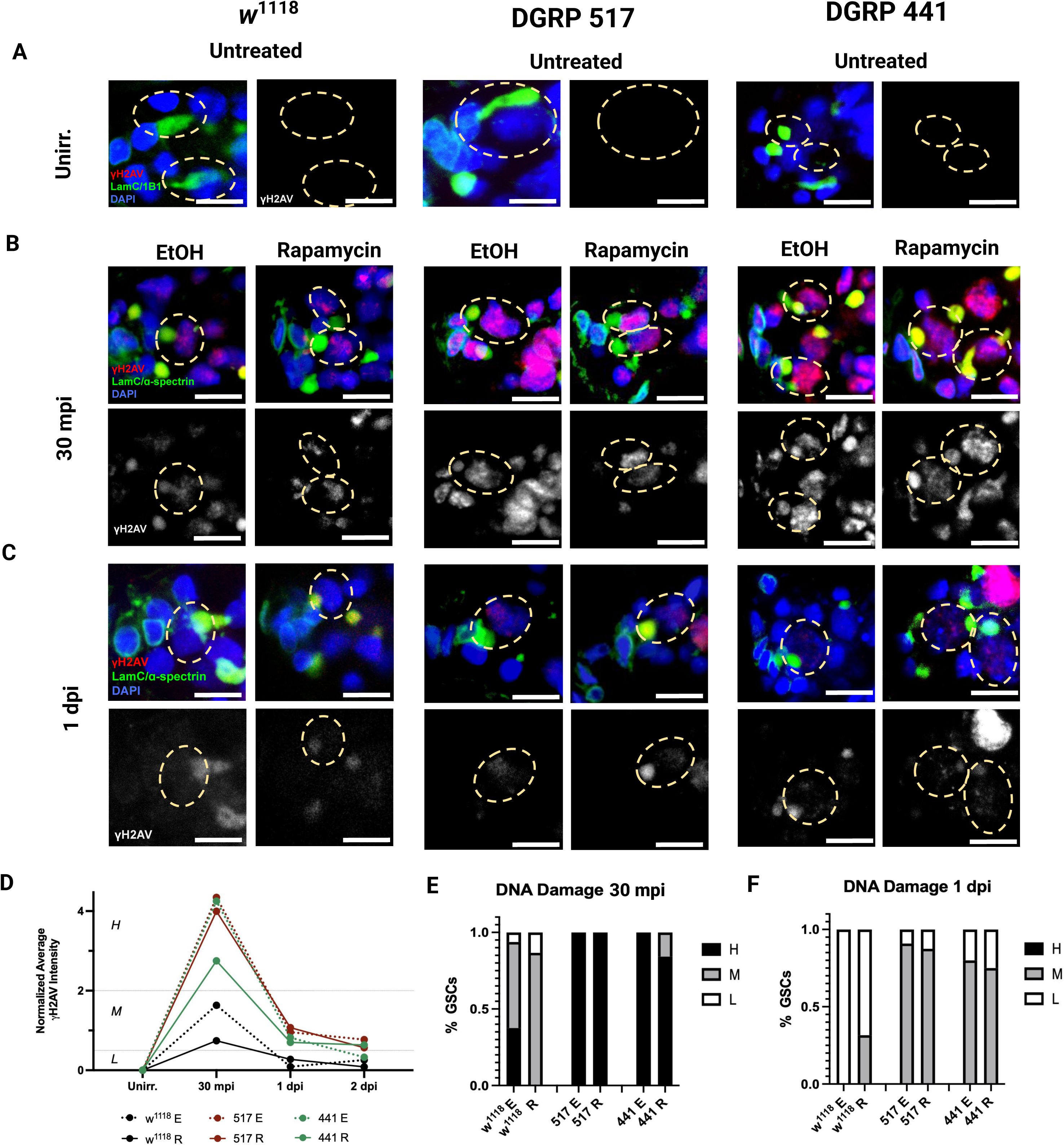
Rapamycin does not impair DNA-DSB repair in GSCs post-IR. (**A**-**C**) Representative confocal images depicting nuclei of GSCs from unirradiated **(A)**, 30 mpi **(B)**, and 1 dpi **(C)** in w^1118^, DGRP 517, and DGRP 441 lines treated with either solvent control (5% EtOH) or rapamycin (200 µM). Stained with LamC (green, CpC and TF), α-spectrin (green, spectrosomes), γ-H2AV (red, DNA-DSB), and DAPI (blue, nuclei) (Scale bar = 10 µm). **(D)** Average intensity for all samples normalized to the intensity of the unirradiated sample in each respective line for unirradiated, 30 mpi, 1 dpi, and 2 dpi timepoints. Samples listed in the legend are w^1118^, control (dotted black); w^1118^, rapamycin (solid black); DGRP 517, control (dotted red); DGRP 517, rapamycin (solid red); DGRP 441, control (dotted green); DGRP 441, rapamycin (solid green). (**E**-**F**) Stacked bar plots showing percentage of GSCs, from flies treated with rapamycin or solvent control, with low (white), moderate (gray), or high (black) DNA damage 30 mpi **(E)** or 1 dpi **(F)** (n=11 to 19 GSCs in E and F).

Quiescence is marked by the upregulation of autophagy and mitophagy that follows the inhibition mTORC1 in Drosophila GSCs [12, 41–43]. We therefore asked whether sensitive and resistant lines differ in their levels or duration of mitophagy when treated with rapamycin.

Actively proliferating stem cells often feature a tubular mitochondrial network localized to the cell anterior [12, 44–46]. In contrast, the fragmented mitochondrial network of quiescent cells is diffused over a larger part of the cell [12, 44, 47]. We assayed for mitophagy by quantifying the number of GSCs with diffuse localization of the mitochondrial marker anti-ATPSynβ (Methods).

In control-treated w^1118^ flies, the percentage of GSCs with fragmented mitochondria increased from 22.7% pre-irradiation to 66.7% at 1 dpi, signifying increased mitophagy following irradiation. At 2 dpi, the percentage dropped to 26.7%, coinciding with quiescence exit (Figure 5D). On the other hand, mitophagy was detected in 52.5% of rapamycin-treated w^1118^ GSC at 2 dpi (Figure 5D). Similar to w^1118^, mitophagy persisted in GSCs from rapamycin-treated DGRP 517 at 2 dpi, however, the frequency of fragmented mitochondria remained high in DGRP 517 from 1 dpi (70.6%) to 2 dpi (75.0%) (Figure 5C, E). Flies of the relatively resistant line DGRP 441 were initially responsive to IR, with the percentage of GSCs with fragmented mitochondria rising to 66.67% in control-treated flies and 69.6% in rapamycin-treated flies at 1 dpi. But at 2 dpi in rapamycin-treated DGRP 441 flies, similar to the control-treated flies, the mitochondrial fragmentation rate dropped at a similar frequency to that in unirradiated flies (Figure 5C, F).

**Figure 5.**
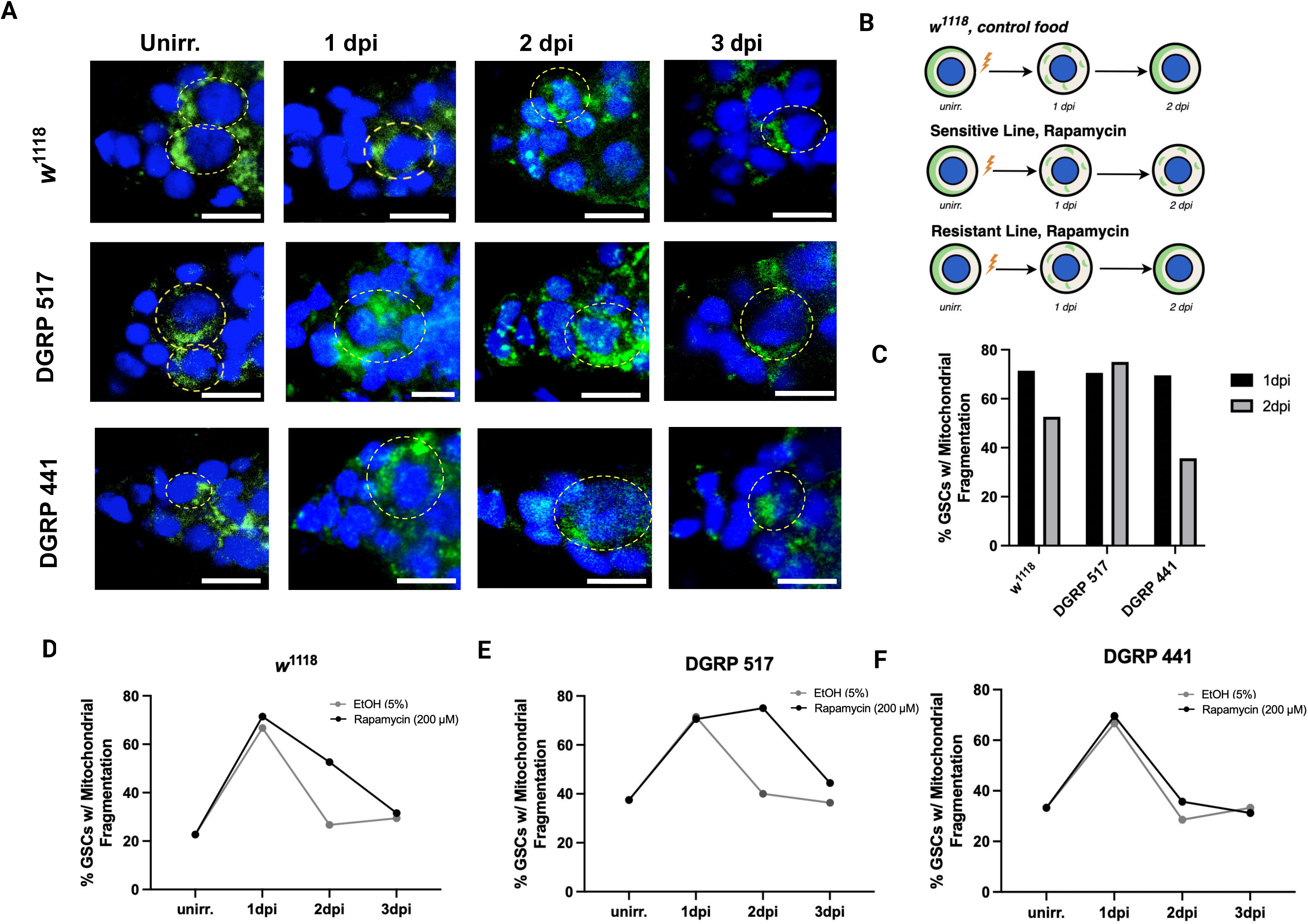
DGRP 517, DGRP 441 and w^1118^ display differential sensitivity to rapamycin in mitophagy. (**A**) Representative confocal microscopy images of GSCs from w^1118^, DGRP 517 and DGRP 441 treated with rapamycin (200 µM) at unirradiated, 1 dpi, 2 dpi, and 3 dpi timepoints. Stained for mitochondria (green, ATP-Synβ) and DAPI (blue, nuclei), (Scale bar = 10 µm) (**B**) Schematic representation of anterior mitochondrial localization and fragmentation in each line treated with rapamycin, in unirradiated, 1 dpi, and 2 dpi timepoints. (**C**) Bar graph representation of the percentage of GSCs with fragmented mitochondria in rapamycin-treated w^1118^, DGRP 517, and DGRP 441 flies at 1 dpi and 2 dpi. (**D**) Percentage of GSCs with fragmented mitochondria in w^1118^ with solvent control or rapamycin treatment across all timepoints (each data point represents an average from n = 15-22 GSCs). (**E**) Percentage of GSCs with fragmented mitochondria in DGRP 517 with solvent control or rapamycin treatment across all timepoints (each data point represents an average from n = 11-18 GSCs). (**F**) Percentage of GSCs with fragmented mitochondria in DGRP 441 with solvent control or rapamycin treatment across all timepoints (each data point represents an average from n = 14-23 GSCs).

Unlike the resolution of DNA damage that followed IR among flies of both the highly sensitive line DGRP 517 and the resistant line DGRP 441 (Figure 4), we find that mitophagy persists concomitantly with the delay in GSC quiescence among these lines. Our results indicate that the duration of quiescence across these genotypes associates with the state of mitophagy and not of DNA damage.

## Discussion

Under genotoxic stress, stem cells enter a transient quiescence to restore cellular integrity before regenerating lost tissue. The inhibition of mTORC1 maintains stem cell quiescence, and its activation promotes cell cycle re-entry. We have shown that the mTORC1-specific inhibitor rapamycin prevents stem cells from exiting quiescence and delays regeneration in the Drosophila female germline. We also show that the effect of rapamycin on quiescence depends on genetic background.

By using fly lines that vary in their sensitivity to rapamycin, we have probed the mechanisms by which the quiescence induced by IR is prolonged by rapamycin. In comparison to the lab strain w^1118^, a sensitive line with more extremely prolonged quiescence, showed concomitantly prolonged mitophagy, whereas in the resistant line with a normal duration of quiescence, mitophagy appeared to resolve quickly (Figure 5). In contrast to the variation in the persistence of mitophagy, we found no evidence for a differential effect of rapamycin on the timing of DNA-DSB repair in w^1118^, compared to DGRP 441 and DGRP 517. Therefore, the genetic variation associated with prolonged quiescence does not appear to affect DNA repair. Taken together, these results suggest that genetic variation in the removal of damaged mitochondria may underlie the range of responses to rapamycin in GSC quiescence among the DGRP. We have previously reported that mTORC1 inhibition, via rapamycin, promotes PINK1/Parkin-mediated mitophagy and mitophagy-dependent degradation of outer-mitochondrially localized G1/S checkpoint-associated cyclin E [12]. The degradation of cyclin E may explain the link between mitophagy and cell cycle arrest [12].

mTORC1 signaling is fundamental to growth and development in a wide range of species, and there are a growing number of cellular and developmental mechanisms that are affected by mTORC1 [48, 49]. The effects of rapamycin therefore may in turn be highly pleiotropic. We have revealed that the genetic variation in the response to rapamycin that appears among the wild-derived DGRP is not consistent across the phenotypes affected by rapamycin (Supplementary Figure S2). Rapamycin both reduces larval size and prolongs the time required for larvae to form pupae [34, 35], and both effects vary strongly by genetic background [30]. By sampling the DGRP lines from those screened for the effect of rapamycin on development time, we were able to test the hypothesis that genetic variation in the effect of rapamycin affects mTORC1 activity generally across phenotypes, such as the pace of development and stem cell quiescence. Consistent with this hypothesis, we found that the effect of rapamycin on quiescence correlates with the rapamycin sensitivity of larval growth among these lines (Figure 2J). In contrast to this hypothesis, however, we found that the extent of developmental delay induced by rapamycin reported in Harrison et al. (2024) did not correlate with the variation in the duration of quiescence that we measured (Supplementary Figure S2B). The difference in the response to rapamycin between genotypes in development therefore appears to affect different mechanisms than those that affect the duration of quiescence. Therefore, rapamycin sensitivity can be defined by line-specific variation in mTORC1 effector pathways, driven by genetic diversity.

Here, we have deciphered that mitophagy underlies variation in rapamycin sensitivity in quiescence. However, further study will be needed to identify common variants across mTORC1 signaling, which will clarify the variation underlying sensitivity and resistance to mTORC1 inhibitors.

## Author Contributions

Conceptualization, S.P., M.G., H.R.B.; methodology, S.P. and M.G.; formal analysis, S.P., B.R.H..; investigation, S.P., M.G., T.Z., Y.G., J.S.; resources, H.R.B.; data curation, S.P., B.R.H.; writing—original draft preparation, S.P.; writing—review and editing, S.P., M.G., H.R.B, B.R.H., D.E.L.P.; visualization, S.P., M.G., T.Z., Y.G.; supervision, S.P., H.R.B.; project administration, S.P., M.G.; funding acquisition, H.R.B., D.E.L.P. All authors have read and agreed to the published version of the manuscript.

## Institutional Review Board Statement

Not applicable.

## Informed Consent Statement

Not applicable.

## Data Availability Statement

Data is contained within the article or Supplementary Materials. The data presented in this study are available in Supplementary Materials.

## Acknowledgments

We thank the members of the Ruohola-Baker lab for their stimulating comments. We thank Christopher Cavanaugh for technical assistance in the irradiation of flies. We thank Dr. Ellen Ward for experimental advice.

## Supplementary Figures

**Supplemental Figure 1.**
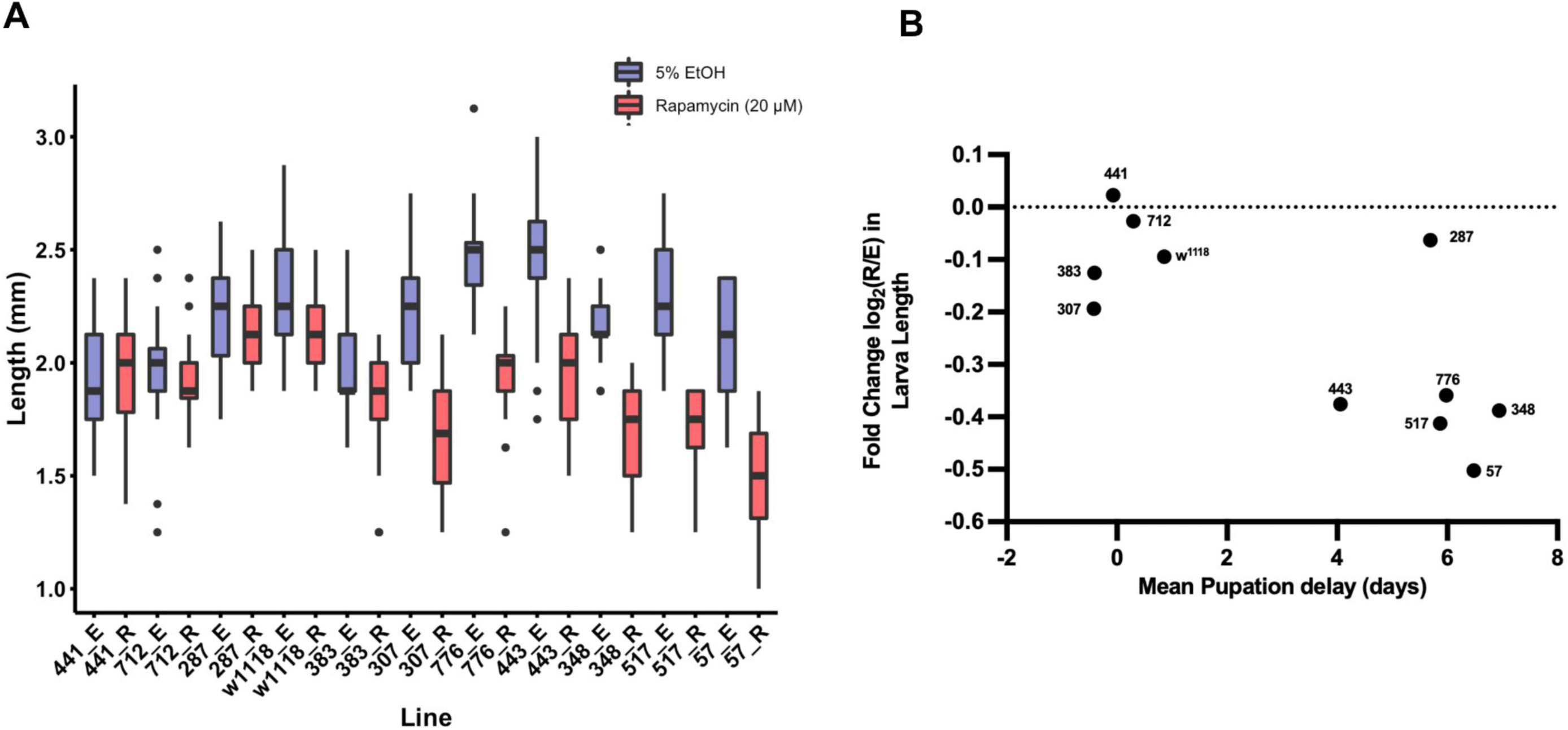
(A) Length (mm) of larvae at 72 h post-oviposition, treated with yeast paste dissolved with 5% EtOH (solvent control) or rapamycin (20 µM) diluted in 5% EtOH (see Methods). Red-shaded box-plots correspond to lengths from rapamycin-treated larvae and blue-shaded box-plots correspond to lengths from control-treated larvae (n = 19-62 larva). Batches were repeated if n<15. (B) Sensitivity to rapamycin in larva length represented as a log_2_ fold change of mean length (rapamycin/control) over the sensitivity to rapamycin in pupation delay (days, Harrison et al., 2024), where Spearman’s correlation (r_s_ = -0.655, P-value = 0.029).

**Supplemental Figure 2.**
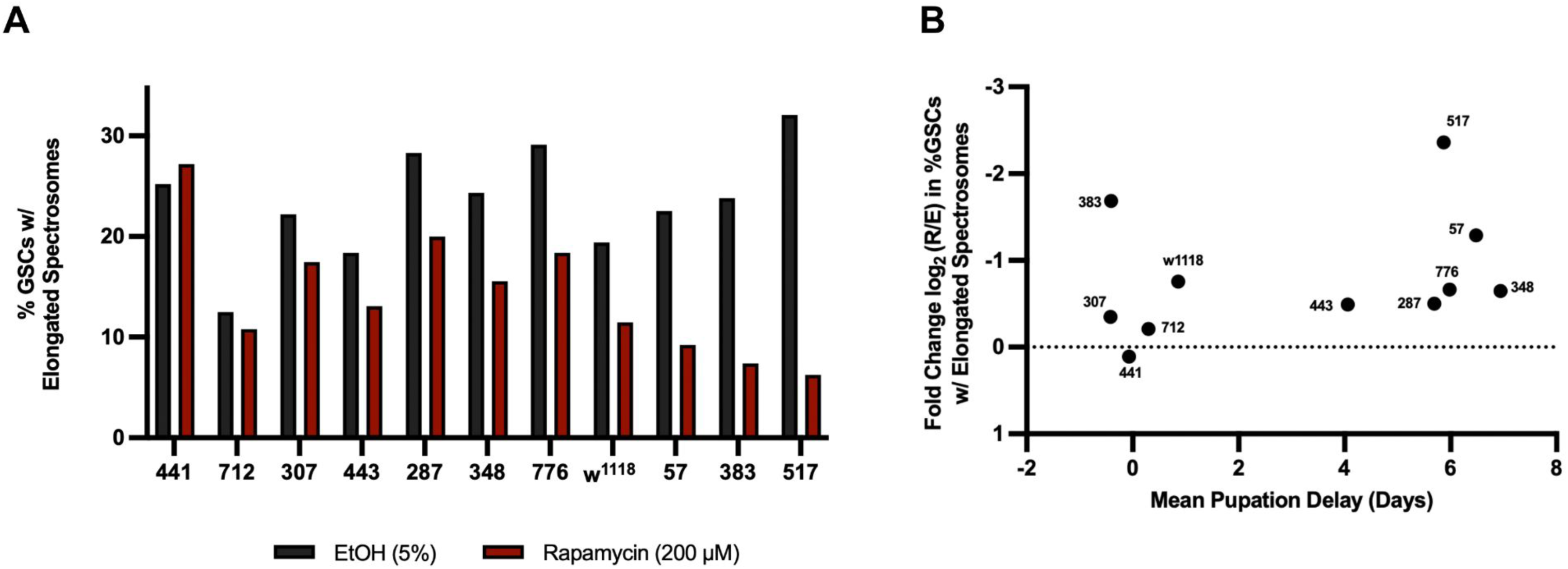
(A) Percentage of GSCs w/ elongated spectrosomes in each DGRP line and *w*^1118^ at 2 dpi treated with rapamycin (200 µM) or 5% EtOH (solvent control). Red corresponds to data for rapamycin-treated flies, and black corresponds to data for control-treated flies (n=158-212 GSCs). (B) The sensitivity to rapamycin in delay in cell-cycle reentry represented as a log_2_ fold change of % GSCs with elongated spectrosomes (rapamycin/control) over the sensitivity to rapamycin in developmental delay (days, Harrison et al., 2024), where Spearman’s correlation (r_s_ = 0.4, P-value = 0.223).

